# Resting-State Connectivity of Auditory and Reward Systems in Alzheimer’s Disease and Mild Cognitive Impairment

**DOI:** 10.1101/2020.03.11.986125

**Authors:** Diana Wang, Alexander Belden, Suzanne Hanser, Maiya R. Geddes, Psyche Loui

**Affiliations:** Harvard College; Northeastern University; Brigham and Women’s Hospital and Harvard Medical School; Berklee College of Music

**Keywords:** resting state fMRI, auditory, reward, dementia, Alzheimer’s disease, mild cognitive mpairment

## Abstract

Music-based interventions have become increasingly widely adopted for dementia and related disorders. Previous research shows that music engages reward-related regions through functional connectivity with the auditory system. Here we characterize intrinsic connectivity of the auditory and reward systems in healthy aging, mild cognitive impairment (MCI) - a predementia phase of cognitive dysfunction, and Alzheimer’s disease (AD). Using resting-state fMRI data from the Alzheimer’s Database Neuroimaging Initiative, we tested functional connectivity within and between auditory and reward systems in older adults with MCI, AD, and age-matched healthy controls (N=105). Seed-based correlations were assessed from regions of interest (ROIs) in the auditory network, i.e. anterior superior temporal gyrus (aSTG), posterior superior temporal gyrus (pSTG), Heschl’s Gyrus, and reward network (i.e., nucleus accumbens, caudate, putamen, and orbitofrontal cortex [OFC]). AD individuals were lower in both within-network and between-network functional connectivity in the auditory network and reward networks compared to MCI and healthy controls. Furthermore, graph theory analyses showed that MCI individuals had higher clustering, local efficiency, degrees, and strengths than both AD individuals and healthy controls. Together, the auditory and reward systems show preserved within- and between-network connectivity in MCI relative to AD. These results suggest that music-based interventions have the potential to make an early difference in individuals with MCI, due to the preservation of functional connectivity in reward-related regions and between auditory and reward networks at that initial stage of neurodegeneration.

## 1 Introduction

Alzheimer’s disease (AD) is a severe and rapidly increasing problem, with over 5 million Americans suffering from this illness. While AD affects 10% of adults over age 65, an additional 15-20% of people above age 65 have mild cognitive impairment (MCI), a predementia phase of cognitive dysfunction linked to higher levels of inflammation and associated with faster clinical deterioration towards dementia (*Alzheimer’s Association: Facts and Figures*, 2019; Pal, et al., 2018; Oikonomidi, et al., 2017; Stella, 2014). In recent years, music-based interventions (MBIs) have become increasingly adopted for patients with Alzheimer’s disease and related disorders. Several randomized controlled trials have shown positive results in the effect of receptive MBIs on alleviating symptoms of cognitive decline, especially in improving mood and reducing stress when listening to familiar music. However, findings to date have been mixed - partly because of variability between subjects, small sample size, and because of differences between intervention protocols across studies (Vink & Hanser, 2018). Part of the challenge in understanding MBIs in neurodegenerative disease is that we do not yet know the influence of cognitive decline on brain networks that are involved in music processing. Advancing this knowledge could help researchers target more precisely when and how to administer MBIs and music therapy.

To date, the best available evidence suggests that music listening may motivate behavior through interactions between brain networks necessary for auditory predictions (such as predictions for melody, harmony, and rhythm) and the brain’s reward system. The auditory cortex is a central hub of an affective-attentional network, home to predictive coding where the brain constructs a hierarchical, generative, top-down model of the world. There is abundant evidence showing that listening to music that we enjoy engages the dopaminergic reward system, indicating that rewarding music has similar properties to other rewarding experiences such as monetary gain and social stimulation (Ferreri et al., 2019; Gold et al., 2019; Salimpoor et al., 2013). When listening to personally pleasurable music, task fMRI has shown that cortical structures in the superior temporal lobe, which constitute an auditory brain network, are correlated in activity with areas in the reward system centering around the ventral striatum (Gold et al., 2019; Martínez-Molina, Mas-Herrero, Rodríguez-Fornells, Zatorre, & Marco-Pallarés, 2016; Salimpoor et al., 2013). Findings from structural neuroimaging have linked white matter connectivity between auditory and reward-related areas, specifically the posterior superior temporal gyrus to the anterior insula and ventromedial prefrontal cortex (vmPFC), to individual differences in reward sensitivity to music (Loui et al., 2017; Martínez-Molina, Mas-Herrero, Rodríguez-Fornells, Zatorre, & Marco-Pallarés, 2019; Sachs, Ellis, Schlaug, & Loui, 2016). These findings suggest that there is a neuroanatomical network that is known to be involved in deriving rewards from music listening (Belfi & Loui, 2019).

In contrast to the structural neuroimaging and task neuroimaging literature, less is known about the intrinsic functional connectivity of the auditory and reward systems, and even less is known about how these patterns of intrinsic functional connectivity may vary in different stages of neurodegeneration. In a landmark study, Jacobsen et al. (2015) compared brain activity of young adults listening to familiar and unfamiliar music in functional Magnetic Resonance Imaging (fMRI), and found that a specific region within the anterior cingulate cortex (ACC) was more active when listening to familiar music, likely part of the auditory prediction network. The authors then analyzed data of essential Alzheimer’s disease biomarkers in a region of interest derived from musical memory findings which included the caudal anterior cingulate cortex and ventral pre-supplementary motor area. They showed that this musical memory region was relatively spared in AD, with minimal cortical atrophy and disruption of glucose-metabolism. These findings support the potential efficacy of MBIs in engaging these relatively preserved brain regions in individuals with AD. Overall, these findings raise the intriguing possibility that music processing might engage brain networks that are relatively spared in neurodegeneration. However, the fMRI results from music listening were obtained from a healthy group of young adults. Thus, results could be explained by intrinsic differences between the different age groups rather than by the specific effects of music per se.

Another study specifically conducted resting state fMRI (rsfMRI) and task fMRI during music listening in the same group of AD patients. King et al. (2019) showed that after listening to familiar music, patients with AD had increased functional connectivity in multiple regions including the default mode network and the auditory networks. While these results provide strong evidence for the use of familiar music in music-based interventions, it remains unclear to what extent these differences in brain connectivity relate to symptom severity in AD and stage of illness. Taken together, it is clear that understanding the intrinsic functional connectivity within and between the auditory and reward systems, and how they change in the aging brain and in different clinical stages of AD, may shed light on how and why music listening could help dementia and promote healthy aging.

The study of intrinsic functional brain networks is aided by recent developments in open science and open data sharing initiatives. The Alzheimer’s Disease Neuroimaging Initiative (ADNI) is a multicenter project that shares neuroimaging data from patients with AD, patients with MCI, and older adult controls (Jack et al., 2008). Data from ADNI offer a starting point from which to investigate intrinsic functional networks at different stages of cognitive decline. The overarching goals of the ADNI study are (1) to detect AD at the earliest possible stage (pre-dementia) and identify ways to track the disease’s progression with biomarkers; (2) to support advances in AD intervention, prevention, and treatment through the application of new diagnostic methods at the earliest possible stages (when intervention may be most effective); and (3) to continually administer ADNI’s innovative data-access policy, which provides all data without embargo to all scientists in the world.

Here we ask how the auditory and reward systems are intrinsically connected in the healthy older adult brain, and how this connectivity changes at different stages of neurodegeneration. We compare resting state networks of three age-matched groups: AD patients, MCI patients, and healthy controls (CN). We identify networks of regions with known roles in auditory prediction and reward, and use them as seed regions of interest to compare the three groups in seed-based connectivity across the brain, whole-brain second-level contrasts to assess between-group differences in resting state functional connectivity, and in ROI-to-ROI connectivity within and across brain networks. Finally, we apply measures from graph theory to describe the complex network properties of the auditory and reward systems, and to see how these networks change in different stages of dementia.

## 2 Materials and Methods

### Sample

We used open-source data from ADNI (Jack et al., 2008). From the available data we limited our sample to patients who had magnetization-prepared, rapid-acquisition, gradient echo (MPRAGE) and rsfMRI scans that were free of artifacts, and that met the specific scan parameters below. This resulted in 105 older adults (ages 55-90) matched in age and gender were selected from the ADNI study set. In the control group (N=47), ages ranged from 56-86, with 27 females; in the MCI group (N=47), ages ranged from 56-88, with 27 females; and in the AD group (N=11), ages ranged from 55-86, with three females. The smaller sample of AD patients is due to lower data quality, because of movement or noise artifacts from the available data. For each individual, two types of data were extracted for use in data analysis: structural MRI (MPRAGE) and functional MRI (functional MRI).

### Procedures

#### MRI Acquisition

High-resolution T1 and resting state images were acquired in a 3T SIEMENS scanner at multiple locations in the United States and Canada. The anatomical images were acquired using a T1-weighted, 3D, MPRAGE volume acquisition with a voxel resolution of 0.8 × 0.8 × 0.8 mm^3^ (TR = 2.3 s, TE = 2.95 ms, flip angle = 9°, Matrix X = 240 pixels, Matrix Y = 256 pixels, Matrix Z = 176 pixels, Mfg Model = Prisma_fit, Pulse Sequence = GR/IR, Slice Thickness = 1.2 mm).

Resting state MRI was acquired as 197 contiguous echo planar imaging (EPI) functional volumes (TR = 3 s; TE = 30 ms; flip angle = 90 degrees; acquisition voxel size = 3.4375 × 3.4375 × 3.4375 mm^3^). Participants kept their eyes open during resting state data acquisition.

#### MRI Preprocessing

Structural and functional MRI preprocessing were carried out with the CONN Toolbox (http://www.nitrc.org/projects/conn) (Whitfield-Gabrieli & Nieto-Castanon, 2012). In order, this consisted of functional realignment and unwarp (subject motion estimation and correction); functional centering to (0,0,0) coordinates (translation); functional slice-timing correction; functional outlier detection (Artifact Detection and Removal Tool (ART)-based identification of outlier scans for scrubbing); functional direct segmentation and normalization (simultaneous grey/white/cerebrospinal fluid segmentation and Montreal Neurological Institute normalization); functional smoothing (spatial convolution with 8 mm Gaussian kernel); structural center to (0,0,0) coordinates (translation); structural segmentation and normalization (simultaneous grey/white/CSF segmentation and MNI normalization). An interleaved slice order was used for Siemens scans, intermediate settings (97^th^ percentiles in normative samples), a global-signal z-value threshold of 9, subject-motion mm threshold of 2, structural target resolution of 1 mm, functionals target resolution of 3.4375 mm, and a bounding box of [90 -126 -72; 90 90 108] mm. Denoising steps for functional connectivity analysis included corrections for confounding effects of white matter and cerebrospinal fluid (Behzadi et al., 2007), and bandpass filtering to 0.008-0.09 Hz.

#### Regions of Interest (ROIs) Selection

When choosing the regions of interest (ROIs) for seed based connectivity measures, we chose regions of interest from the CONN default atlas (Whitfield-Gabrieli & Nieto-Castanon, 2012) which contains 185 ROIs and 32 networks. We included all ROIs in the superior, middle, and inferior temporal lobes, resulting in 18 ROIs: right anterior Superior Temporal Gyrus (aSTGR), left anterior Superior Temporal Gyrus (pSTGR), right posterior Superior Temporal Gyrus (pSTGR), left posterior Superior Temporal Gyrus (pSTGL), right anterior Middle Temporal Gyrus (aMTGR), left anterior Middle Temporal Gyrus (aMTGL), right posterior Middle Temporal Gyrus (pMTGR), left posterior Middle Temporal Gyrus (pMTGL), right temporooccipital Middle Temporal Gyrus (toMTGR), left temporooccipital Middle Temporal Gyrus (toMTGL), right anterior Inferior Temporal Gyrus (aITGR), left anterior Inferior Temporal Gyrus (aITGL), right posterior Inferior Temporal Gyrus (pITGR), left posterior Inferior Temporal Gyrus (pITGL), right temporooccipital Inferior Temporal Gyrus (toITGR), left temporooccipital Inferior Temporal Gyrus (toITGL), right Heschl’s Gyrus (HGR), and left Heschl’s Gyrus (HGL).

Then, we selected 18 ROIs as valuation and reward-related regions based on the previous literature (Belfi & Loui, 2019): Right Insular Cortex (InsulaR), Left Insular Cortex (InsulaL), Anterior Cingulate Gyrus (AC), Posterior Cingulate Gyrus (PC), right Frontal Orbital Cortex (FOrbR), left Frontal Orbital Cortex (FOrbL), right Caudate (CaudateR), left Caudate (CaudateL), right Putamen (PutamenR), left Putamen (PutamenL), right Pallidum (PallidumR), left Pallidum (PallidumL), right Hippocampus (HippocampusR), left Hippocampus (HippocampusL), right Amygdala (AmygdalaR), left Amygdala (AmygdalaL), right Accumbens (AccumbensR), left Accumbens (AccumbensL).

Finally, we combined the 18 auditory ROIs together into an *Auditory Network* and the 18 reward ROIs together into a Reward/Valuation Network (hereafter *Reward Network*). Figure 1 shows the auditory and reward network ROIs.

**Figure 1.**
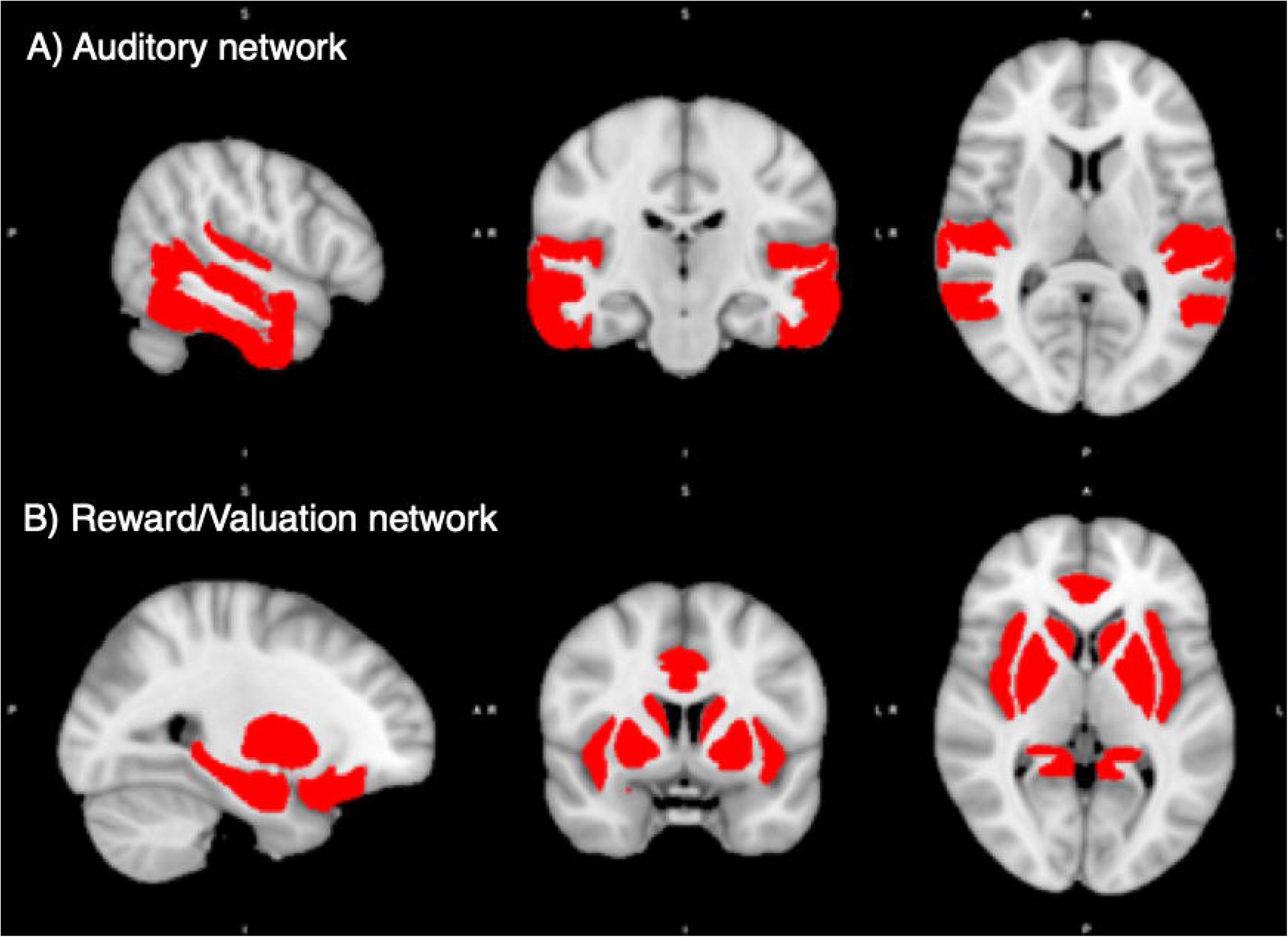
Regions of interest in the auditory and reward/valuation networks from the CONN Toolbox. A: compilation of the 18 auditory ROIs from CONN. B: compilation of the 18 reward ROIs from CONN. See Table 1 for a list of the ROIs used.

### Seed-Based Connectivity Analyses

Since we were interested in whole-brain connectivity patterns of the auditory and reward networks, we first seeded the auditory and reward networks defined above, and for each group of subjects we extracted all voxels that were significantly functionally connected (using bivariate correlation) to the seed ROIs at the p<0.05, p-Family Wise Error corrected level. Slices were chosen at the peak cluster for all three groups. Between-group comparisons within our auditory and reward networks were additionally conducted where we extracted all voxels that were significantly functionally connected to the seed ROIs at the p<0.05, p-Family Discovery Rate cluster-size corrected level.

**Table 1:** Auditory and reward brain regions and the XYZ-coordinates of their centers of gravity. The 36 ROI’s are from the default atlas in the CONN Toolbox (Whitfield-Gabrieli and Nieto-Castanon, 2012). Coordinates in millimeters in the Montreal Neurological Institute space.

| Seed | Side | Coordinates |  |  |
| --- | --- | --- | --- | --- |
|  |  | x | y | z |
| <u>Auditory Network</u> |  |  |  |  |
| Anterior Superior Temporal Gyrus | Left | -56.17233662 | -3.906445837 | -7.970008953 |
|  | Right | 57.50133452 | -0.763345196 | -10.16592527 |
| Posterior Superior Temporal Gyrus | Left | -62.28850325 | -29.1691974 | 3.79640533 |
|  | Right | 61.34069304 | -23.9858939 | 1.573443729 |
| Anterior Middle Temporal Gyrus | Left | -57.46777317 | -4.205255878 | -22.13914246 |
|  | Right | 57.88912197 | -1.522025772 | -24.50584357 |
| Posterior Middle Temporal Gyrus | Left | -60.90641565 | -27.35880095 | -10.99670993 |
|  | Right | 61.07506629 | -22.52454969 | -12.14985828 |
| Temporooccipital Middle Temporal Gyrus | Left | -57.63985911 | -52.99985324 | 0.8241855 |
|  | Right | 58.18120733 | -49.22239081 | 1.59748564 |
| Anterior Inferior Temporal Gyrus | Left | -48.14201402 | -4.974548137 | -39.1910734 |
|  | Right | 46.22887061 | -2.40987285 | -41.10583396 |
| Posterior Inferior Temporal Gyrus | Left | -53.44317202 | -28.4572097 | -25.98756311 |
|  | Right | 53.42195698 | -23.46366737 | -28.13352571 |
| Temporooccipital Inferior Temporal Gyrus | Left | -51.81781305 | -53.44356261 | -16.53315697 |
|  | Right | 54.14167857 | -49.87783595 | -16.73076313 |
| Heschl's Gyrus | Left | -45.19664938 | -20.3214998 | 7.192261667 |
|  | Right | 46.11156095 | -17.40440846 | 6.967611336 |
| <u>Reward Network</u> |  |  |  |  |
| Insular Cortex | Left | -36.3943 | 1.1868 | 0.0824 |
|  | Right | 37.3847 | 2.5497 | -0.1738 |
| Anterior Cingulate Gyrus | Bilateral | 0.80277298 | 18.29370562 | 24.34508732 |
| Posterior Cingulate Gyrus | Bilateral | 0.784845018 | -36.62174953 | 29.97508841 |
| Frontal Orbital Cortex | Left | -29.54284237 | 23.66228394 | -16.57261043 |
|  | Right | 29.11395129 | 23.07066013 | -16.23143128 |
| Caudate | Left | -12.78549849 | 8.976992796 | 9.737392517 |
|  | Right | 13.30156062 | 10.01080432 | 10.49051621 |
| Putamen | Left | -24.90149125 | 0.482649842 | 0.339546888 |
|  | Right | 25.49574896 | 1.776008657 | 0.30344721 |
| Pallidum | Left | -18.95779356 | -5.120351024 | -1.333890514 |
|  | Right | 19.85048905 | -4.005123428 | -1.189101071 |
| Hippocampus | Left | -25.17773788 | -23.1916109 | -13.80594092 |
|  | Right | 26.49706667 | -20.95893333 | -14.25013333 |
| Amygdala | Left | -22.99501151 | -4.94666155 | -17.73177283 |
|  | Right | 23.08748616 | -3.985234404 | -17.68696936 |
| Accumbens | Left | -9.463882619 | 11.496614 | -7.170428894 |
|  | Right | 9.368263473 | 12.20359281 | -6.534431138 |

### ROI-to-ROI Analyses and Graph Theory Analyses

R-correlation values for each of the 36 regions of interest from the CONN atlas were extracted for every participant and averaged across each group to compute pairwise correlations and graph theory analyses. Correlation matrices comparing all 36 regions of interest from the CONN atlas were extracted for each participant in each group. These matrices were then exported into Matrix Laboratory (MATLAB) and analyzed using the Brain Connectivity Toolbox (Rubinov & Sporns, 2010). Each network statistic was computed at a range of correlation thresholds from r = 0.05 to r = 0.5. Individual participants’ measures of degrees, clustering coefficients, strengths, betweenness centrality, and local efficiency were calculated for each region in each brain and then averaged across participants for each group, whereas modularity was a single measure for the whole brain that was calculated for each participant. These group averages were then compared using one-way ANOVAs to determine group differences in each network measure while correcting for false-discovery rate of 0.05 for comparisons across 6 network measures (Benjamini & Hochberg, 1995).

## 3 Results

### Seed-Based Connectivity Analyses

Seed-based connectivity patterns for each group are shown in Figure 2. All groups showed highly significant auditory network functional connectivity to the auditory areas, including the STG, MTG, and ITG, at the p < .05 FWE-corrected level. The control and MCI groups additionally showed significant functional connectivity in the parietal, occipital, and frontal lobes. The AD group showed less significant functional connectivity than the other two groups, with the significant functional connectivity only observed in the temporal lobe, and not in the other lobes. Between-group comparisons showed higher functional connectivity in the precuneus for the control group compared to the AD group (p < .05 cluster-size FDR-corrected level). No other between-group differences survived correction for multiple comparisons in seed-based connectivity.

**Figure 2:**
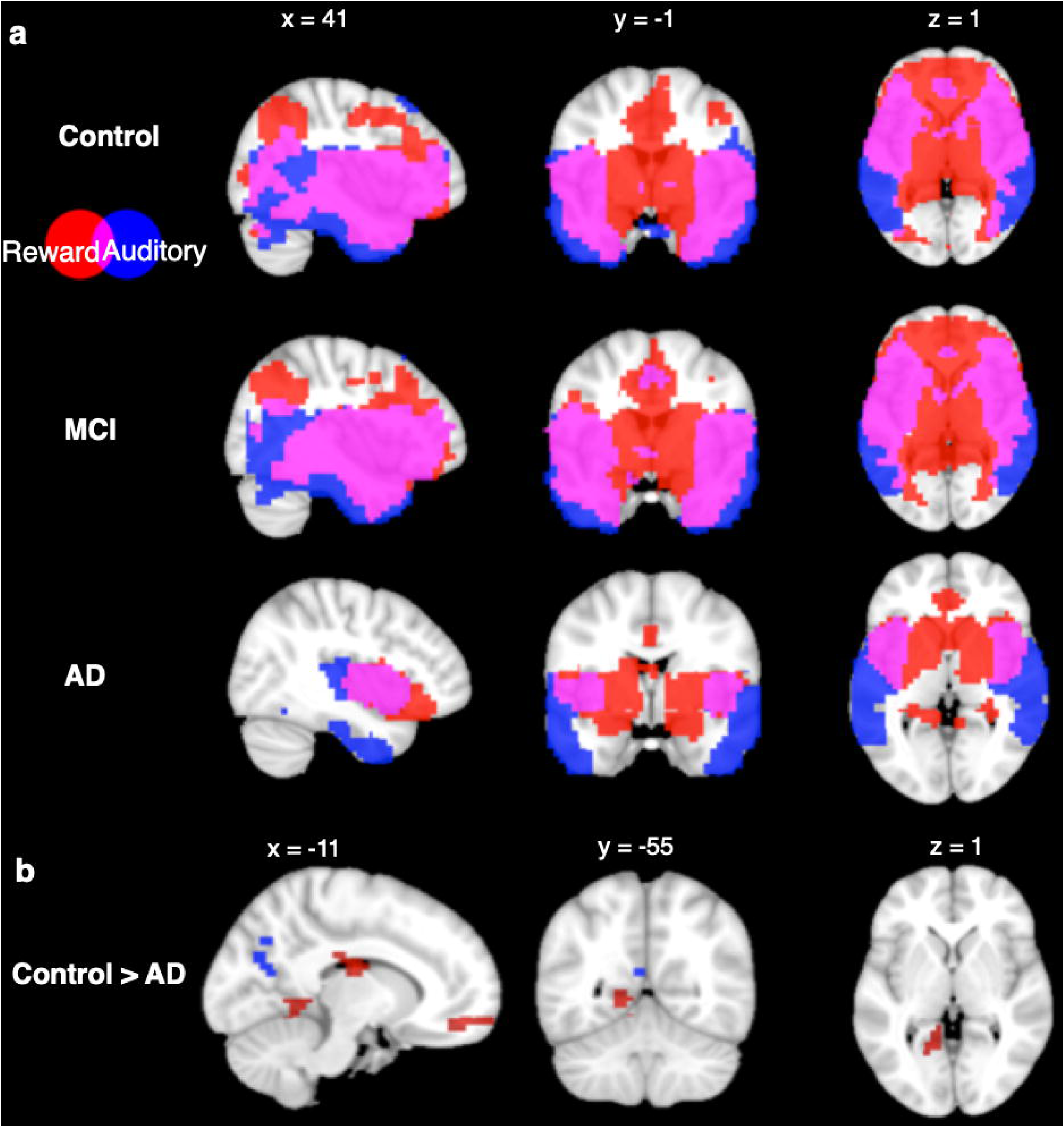
Seed Based Connectivity Analysis. **a**: Connectivity profiles of Control group (top row), MCI group (middle row), and AD group (bottom) for seed regions in the auditory (blue) and reward (red) networks (*p* < 0.05, voxel-wise p-FWE corrected). **b**: Connectivity profile differences comparing Control and AD groups seeded from auditory (blue) and reward (red) networks (*p* < 0.05, p-FDR cluster-size corrected).

Seed-based connectivity from the reward network showed significant functional connectivity within areas of the reward network in all groups at the p < .05 FWE-corrected level. CN and MCI groups both have significant functional connectivity to the auditory network ROIs including the MTG and ITG, as well as significant overlap between areas that are functionally connected to auditory and reward ROIs in the frontal, parietal, and occipital lobes. In contrast, the AD group did not show connectivity in lateral frontal, parietal, or occipital lobes from the reward network ROIs. Between-group comparisons showed higher functional connectivity in the control group compared to the AD group at the p < .05 cluster-size FDR-corrected level in six regions: the cingulate cortex, the medial prefrontal cortex, the left lingual gyrus, the bilateral fusiform gyri, and superior parietal lobule. No other between-group differences were significant.

### ROI-to-ROI Analyses

We further characterized within- and between-network connectivity across the 36 ROIs from the defined auditory and reward networks. Figure 3 shows t-maps of bivariate correlations between each pair of ROIs in each group. All three groups show higher connectivity within each network (auditory-auditory, reward-reward) than between networks (auditory-reward), as shown by higher T values within the diagonal quadrants (which represent auditory-auditory and reward-reward connectivity) than in the off-diagonal quadrants (which represents auditory-reward connectivity). The t-values are generally similar between CN and MCI groups. In contrast, the AD group has lower network connectivity overall.

**Figure 3:**
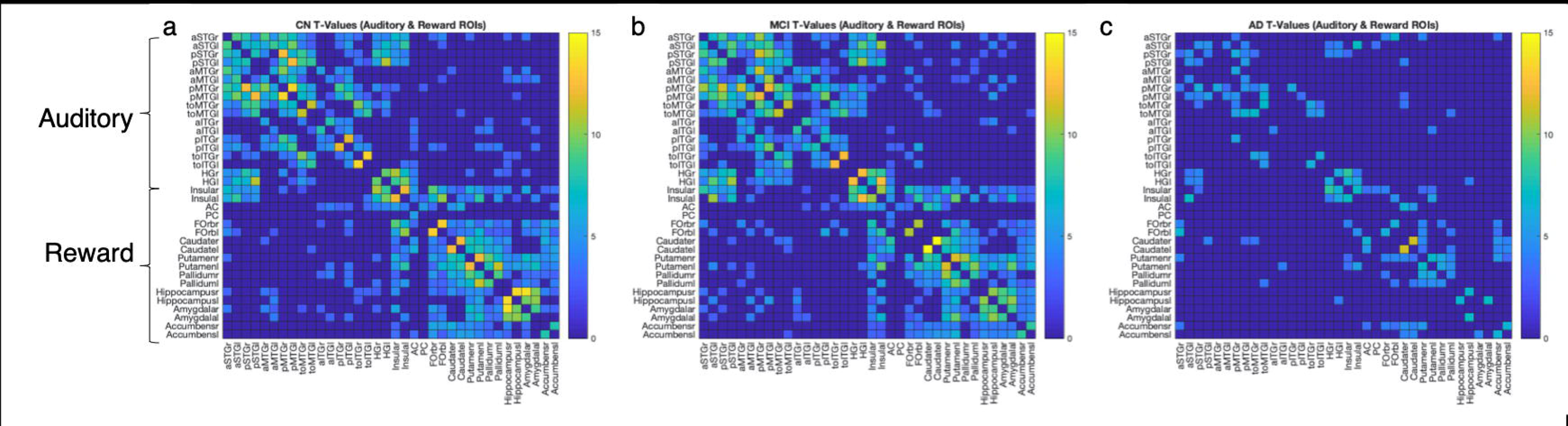
ROI-to-ROI connection matrices and corresponding brain connectomes. a: Control group, b: MCI group, c: AD group showing significant positive correlations (*p* < 0.05, p-FDR corrected) between the auditory and reward regions. The colors correspond to strength of correlation between two ROIs.

### Graph Theory Analyses

Betweenness centrality, degrees, and strengths showed significant main effects of group at a correlation threshold of r = 0.2. Network measures clustering and local efficiency did not have significant main effects of group at this correlation threshold. *Betweenness centrality*, which is the number of shortest paths from one node to another that contains a given node, showed highest levels in the CN group, followed by MCI and then by AD (F(2, 105) = 6.64, p = 0.0019, Benjamini-Hochberg corrected, Figure 4a). On the other hand, the network measure of *degrees*, or the number of nodes significantly correlated to a given node, showed a significant main effect of group as well (F(2, 105) = 4.75, p = 0.0106, Benjamini-Hochberg corrected, Figure 4b), although this time the measure showed highest levels in MCI individuals, followed by CN group, and trailed by the AD group. We see a significant difference between the MCI and AD groups here in which the MCI group has a higher degrees parameter. A similar pattern was also seen in *strengths*, the sum of the correlation coefficients for a given node, with highest levels once again being seen in the MCI group (F(2, 105) = 3.88, p = 0.0237, Benjamini-Hochberg corrected, Figure 4e), and also having a significant difference between the MCI and AD groups, with the MCI group having highest strength. *Clustering coefficient*, the fraction of nodes correlated with a given node that are also correlated with one another, did not show a main effect of group, but was highest in the MCI group, then AD group, and then CN group (F(2, 105) = 1.66, p = 0.1954, Benjamini-Hochberg corrected, Figure 4b). Here, the CN and MCI groups are both significantly higher than the AD groups. *Local efficiency*, the average connectedness in the neighborhood of a given node, also did not show a main effect of group, with the same relative pattern of MCI to AD to CN group (F(2, 105) = 1.64, p = 0.1994, Benjamini-Hochberg corrected, Figure 4c).

**Figure 4:**
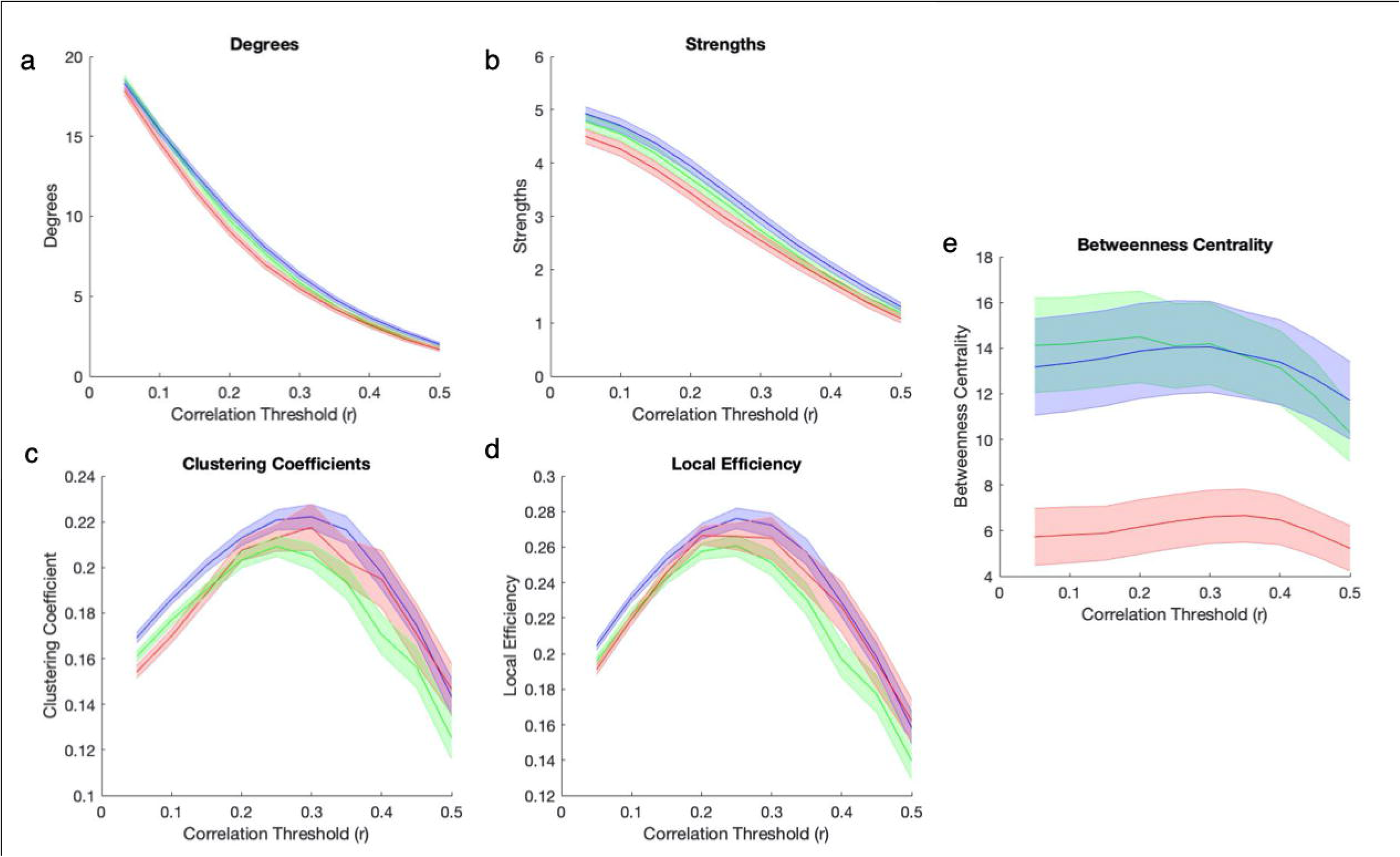
Group Differences in Small World Brain Connectivity. Network measures of degree (a), strengths (b), clustering coefficient (c), local efficiency (d), and betweenness centrality (e) for Control group (green), MCI group (blue), and AD group (red) across a range of correlation thresholds (solid line = mean of all subjects/ROI’s for each group, error bar = standard error for all 36 CONN ROI’s averaged across subjects for each group).

Taken altogether, the MCI group is higher than the CN group in degrees, strengths, clustering, and local efficiency, and is indistinguishable from the CN group in betweenness centrality. The AD group is indistinguishable from others in clustering or local efficiency, while being lower than MCI and CN groups in degrees and strengths, and much lower than both other groups in betweenness centrality. In summary, the pattern of graph theory results show that MCI individuals have consistently high between-network connections as well as within-network clustering within the reward network relative to controls and AD individuals.

## 4 Discussion

Although abundant research supports the interaction between auditory and reward systems in enabling pleasure in music listening, little is known about the intrinsic functional connectivity between the auditory and reward systems. Here, we defined an auditory network and a reward network based on previous studies, and characterized their intrinsic functional connectivity using resting state fMRI from a large sample of AD, MCI, and age-matched controls. We found decreased functional connectivity within and between the two systems in AD individuals. These differences are observable in seed-based as well as ROI-to-ROI connectivity, and also in disruptions that affect degrees, strengths, and betweenness centrality of the overall network.

Importantly, we observe an overlap between seed-based connectivity patterns from the auditory network and the reward network. This overlap was observed in all three groups, centering around the anterior insula. Importantly, there was no overlap among the ROIs chosen as the seed regions of the auditory and reward networks; thus the results are due to similar patterns in functional connectivity between the anterior insula and both the auditory and reward regions. The anterior insula is part of the salience network, which has been posited as a hub that enables alternating between default mode and executive control networks (Menon & Uddin, 2010). The present results extend that previous work, by suggesting that the salience network, with anterior insula at its core, may be key to interactions between large-scale brain systems more generally. This result has important implications. First, it supports the neuroanatomical model for the reward of music listening and music-based interventions, as laid out in Belfi & Loui (2019), which posits that anterior insula is connected to both auditory and reward systems. This finding is also consistent with lesion mapping studies: Cases of acquired musical anhedonia (i.e. the lack of emotional responses to music due to brain injury) mostly have lesions in the anterior insula (Griffiths et al, 2004; Satoh et al, 2011). Thus, the anterior insula seems to be a key region for deriving reward from music listening.

Relative to AD individuals, MCI individuals show preserved functional connectivity, with no significant between-group differences in auditory-seeded or reward-seeded connectivity patterns from age-matched controls. Graph theory results showed higher degrees, strengths, clustering, and local efficiency in the MCI group than in both the AD and the control groups. Thus, the relationship between dementia severity and network connectedness appears to follow an inverse u-shaped curve, with the slightly impaired MCI group showing the strongest and most efficient connections across all the ROIs of the auditory and reward networks. This is different from previous findings in graph theory analysis of resting state networks of MCI, AD and CN groups (Seo et al., 2013). Using FDG-PET data, previous work has shown lower clustering in both MCI and AD groups compared to the CN group. However, those with very mild AD had lower clustering compared to those with mild AD (Seo et al., 2013). On the other hand, a more recent study found that the small world index, a summary network statistic, was significantly decreased in MCI converters who progressed to AD, compared to stable MCI individuals who did not progress to AD (Miraglia et al, 2020). Taken together, the distinctions between MCI and AD may be more fine-grained than are captured in our study. Furthermore, as we were specifically interested in the auditory and reward networks, we used only a subset of ROIs that represented these networks, rather than ROIs covering the whole brain. Thus, our results should not be interpreted as generalizable towards the whole brain in all MCI individuals, but rather as results of a specific hypothesized network of regions important for deriving rewards from music listening.

In the present study, the finding of higher network statistics in auditory and reward network ROIs among MCI individuals may suggest that auditory and reward regions more readily connect to each other in the MCI brain. This may have important implications for music therapy. As music-based interventions rely on the participants’ engagement with music, and the activity and connectivity of the reward system is reflective of engagement in music and in other domains (Ferreri et al., 2019; Kampe, Frith, Dolan, & Frith, 2001; Martínez-Molina et al., 2016; Salimpoor et al., 2013; Tamir & Mitchell, 2012), the current results may suggest that targeting individuals with MCI can capitalize on the heightened auditory-reward connectivity in MCI, thus offering the best chance for effective intervention.

Although AD individuals have less functional connectivity overall, they still show some preserved overlap between auditory and reward systems in the anterior insula. This finding may also have implications for music-based interventions. Specifically, it may be possible to identify specific experiences that also engage the insula, and tailor music-based interventions to maximize these experiences. For example, the anterior insula has been implicated in specificity for voice processing, and has been described as part of a voice-selective cortex (Abrams et al., 2013). Perhaps listening to music with the voice, or even engaging in vocalization in an active music-based intervention, may be specific ways to tap into the reward system. Since the dopaminergic reward system is crucial for motivated behavior, understanding its connectivity patterns to the rest of the brain, and in different stages of disease, offers insight into the design of effective interventions for diseases and disorders.

## 5 Conclusion

We have identified an anatomical model of auditory and reward systems, and characterized the functional connectivity within and between these systems in healthy older adults and in older adults with MCI and AD. Results inform music-based interventions by highlighting the importance of focusing on the MCI population, as they have the most functional connectivity in their auditory and reward systems.

## 6 Author Contributions

PL conceptualized the idea behind this manuscript. DW acquired and preprocessed the behavioral and neuroimaging data, performed data analyses, and wrote the first draft. AB, SH, and MG provided feedback, guidance and support on conceptual and technical aspects of the study. All authors revised the manuscript and approved the submission.

## 8 Acknowledgments

Supported by the Grammy Foundation, NSF-CAREER #1945436, and Kim & Glen Campbell Foundation to PL and the Sidney Baer Foundation Clinical Research Grant to MRG.

## References

Abrams, D. A., Lynch, C. J., Cheng, K. M., Phillips, J., Supekar, K., Ryali, S., … Menon, V. (2013). Underconnectivity between voice-selective cortex and reward circuitry in children with autism. Proc Natl Acad Sci U S A, 110 (29), 12060–12065.

Belfi, A., & Loui, P. (2019). Musical anhedonia and rewards of music listening: Current advances and a proposed model. Annals of the New York Academy of Sciences, In press.

Benjamini, Y., & Hochberg, Y. (1995). Controlling the false discovery rate: a practical and powerful approach to multiple testing. Journal of the royal statistical society. Series B (Methodological), 289–300.

Ferreri, L., Mas-Herrero, E., Zatorre, R. J., Ripollés, P., Gomez-Andres, A., Alicart, H., … Rodriguez-Fornells, A. (2019). Dopamine modulates the reward experiences elicited by music. Proceedings of the National Academy of Sciences, 201811878.

Gold, B. P., Mas-Herrero, E., Zeighami, Y., Benovoy, M., Dagher, A., & Zatorre, R. J. (2019). Musical reward prediction errors engage the nucleus accumbens and motivate learning. Proc Natl Acad Sci U S A.

Jack, C. R., Bernstein, M. A., Fox, N. C., Thompson, P., Alexander, G., Harvey, D., … Weiner, M. W. (2008). The Alzheimer’s disease neuroimaging initiative (ADNI): MRI methods. Journal of Magnetic Resonance Imaging, 27 (4), 685–691.

Jacobsen, J.-H., Stelzer, J., Fritz, T. H., Chételat, G., La Joie, R., & Turner, R. (2015). Why musical memory can be preserved in advanced Alzheimer’s disease. Brain.

Kampe, K. K., Frith, C. D., Dolan, R. J., & Frith, U. (2001). Reward value of attractiveness and gaze. Nature, 413 (6856), 589.

King, J. B., Jones, K. G., Goldberg, E., Rollins, M., MacNamee, K., Moffit, C., … Foster, N. L. (2019). Increased Functional Connectivity After Listening to Favored Music in Adults With Alzheimer Dementia. J Prev Alzheimers Dis, 6 (1), 56–62.

Loui, P., Patterson, S., Sachs, M. E., Leung, Y., Zeng, T., & Przysinda, E. (2017). White Matter Correlates of Musical Anhedonia: Implications for Evolution of Music. Frontiers in Psychology, 8(1664).

Martínez-Molina, N., Mas-Herrero, E., Rodríguez-Fornells, A., Zatorre, R. J., & Marco-Pallarés, J. (2016). Neural correlates of specific musical anhedonia. Proceedings of the National Academy of Sciences, 113 (46), E7337–E7345.

Martínez-Molina, N., Mas-Herrero, E., Rodríguez-Fornells, A., Zatorre, R. J., & Marco-Pallarés, J. (2019). White Matter Microstructure Reflects Individual Differences in Music Reward Sensitivity. The Journal of Neuroscience, 39 (25), 5018.

Menon, V., & Uddin, L. Q. (2010). Saliency, switching, attention and control: a network model of insula function. Brain Structure and Function, 214 (5-6), 655–667.

Oikonomidi, A. et al. (2017). Macrophage Migration Inhibitory Factor is Associated with Biomarkers of Alzheimer’s Disease Pathology and Predicts Cognitive Decline in Mild Cognitive Impairment and Mild Dementia. Journal of Alzheimer’s disease: JAD 60, 273–281, doi: 10.3233/jad-170335

Pal, K., Mukadam, N., Petersen, I. & Cooper, C. (2018). Mild cognitive impairment and progression to dementia in people with diabetes, prediabetes and metabolic syndrome: a systematic review and meta-analysis. Social psychiatry and psychiatric epidemiology 53, 1149–1160, doi: 10.1007/s00127-018-1581-3

Rubinov, M., & Sporns, O. (2010). Complex network measures of brain connectivity: Uses and interpretations. Neuroimage, 52 (3), 1059–1069.

Sachs, M. E., Ellis, R. J., Schlaug, G., & Loui, P. (2016). Brain Connectivity Reflects Human Aesthetic Responses to Music. Social, Cognitive, and Affective Neuroscience, 11 (6), 884–891.

Salimpoor, V. N., van den Bosch, I., Kovacevic, N., McIntosh, A. R., Dagher, A., & Zatorre, R. J. (2013). Interactions between the nucleus accumbens and auditory cortices predict music reward value. Science, 340 (6129), 216–219.

Seo, E. H., Lee, D. Y., Lee, J.-M., Park, J.-S., Sohn, B. K., Lee, D. S., … Woo, J. I. (2013). Whole-brain Functional Networks in Cognitively Normal, Mild Cognitive Impairment, and Alzheimer’s Disease. PLOS ONE, 8 (1), e53922.

Stella, F. et al. (2014). Neurobiological correlates of apathy in Alzheimer’s disease and mild cognitive impairment: a critical review. Journal of Alzheimer’s disease: JAD 39, 633–648, doi: 10.3233/jad-131385

Tamir, D. I., & Mitchell, J. P. (2012). Disclosing information about the self is intrinsically rewarding. Proc Natl Acad Sci U S A, 109 (21), 8038–8043.

Vink, A., & Hanser, S. (2018). Music-Based Therapeutic Interventions for People with Dementia: A Mini-Review. Medicines, 5 (4).

Whitfield-Gabrieli, S., & Nieto-Castanon, A. (2012). Conn: a functional connectivity toolbox for correlated and anticorrelated brain networks. Brain Connect, 2 (3), 125–141.

